## Supplemental Tables 1 t0 3 and captions for supplemental figures and files for "Mealtime alters daily rhythm in nuclear O-GlcNAc proteome to regulate hepatic gene expression"

**S1 Table. Comparison between our study and published liver phosphoproteome datasets.**

| Reference | Number of phospho-peptides | Number of phospho-proteins | Number of oscillating phospho-peptides (Percentage) | Number of oscillating phospho-proteins (Percentage) | Circadian kinases | Normalize to protein level | Label method | TRF | LD vs DD | Cell fraction | Time points X days | Replicate number (total sample number) |
| --- | --- | --- | --- | --- | --- | --- | --- | --- | --- | --- | --- | --- |
| This study | 19593 | 3272 | 1054 (5.38%) | 579 (17.70%) | AKT, S6K, CDK, VRK | Yes | TMT | Yes (3 weeks) | LD | Nucleus | 6 time-points X 1 day | 3 (18) |
| Robles et al. [2] | 7896 | 2672 | 2123 (26.89%) | 1088 (40.71%) | ERK, MEK, RSK, JNK, AKT, mTOR, p70S6K Kinase,  CK1δ | No | No | No | DD | Whole cell | 8 time-points X 2 days | 4 (64) |
| Huang et al. [4] | NA | 2639 | NA | 1668 (63.21%) | NA | No | No | Yes (1 week) | LD | Whole cell | 6 time-points X 2 day | 4 (48) |
| Wang et al. [5] | 9465 | 1657 | NA | 89 (5.37%) | NA | No | No | No | LD | Whole cell | 8 time-points X 2 days | 1 (Pooled 3 mice) (16) |
| Wang et al. [5] | 1448 | NA | 154 (10.64%) | 113 | GSK3α, GSK3β,CK1α, CK1δ, CDK1, CDK4, CDK6 | No | SILAC | Yes (4 days) | LD | Nucleus | 8 time-points X 2 days | 1 (16) |

**S2 Table. Known functions of rhythmic phosphorylation sites that are identified in protein complexes.**

| Gene name | Uniprot ID | Phosphosite | Kinase | Function | Reference |
| --- | --- | --- | --- | --- | --- |
| *Hdac1* | O09106 | S421, S423 | CK2 | Phosphorylation of S421 and S423 promotes the deacetylase activity and complex formation of HDAC1 | [6] |
| *Clock* | O08785 | T461 | CDK5 | T461 phosphorylation reduces stability and promotes nuclear translocation of CLOCK | [7] |
|  |  | S431 | NA | Phosphorylation of S431 primes the phosphorylation of S427 | [8] |
|  |  | S427 | GSK3β | Phosphorylation of S427 leads to CLOCK degradation | [8] |
|  |  | S845 | AKT | Phosphorylation of S845 inhibits its nuclear translocation and affects circadian gene expression | [9] |
| *Rbl2* | Q64700 | S410, S659 | NA | Phosphorylation of S410 and S659 (S413 and S662 in human homolog) promotes the dissociation between RBL2 and E2F4, and relieves repression of E2F4 target gene expression. | [10] |
| *Sfpq* | Q8VIJ6 | T679 | GSK3 | Phosphorylation of T679 (T687 in human homolog) increases the interaction between SFPQ and TRAP150, which inhibits the binding of SFPQ pre-mRNA and thus affects its RNA splicing activity. | [11] |

**S3 Table. The domains within or nearby of the rhythmic phosphosites with unknown function, and only phosphosites identified in protein complexes are included.**

| Gene name | Uniprot ID | Phosphosite | Nearby domains* (amino acid number of the domains) | Distance from nearby domain |
| --- | --- | --- | --- | --- |
| *Ptbp2* | Q91Z31 | S308 | RNA recognition motif (aa309-434) | -1 |
| *Hnrnpm* | Q9D0E1 | S636 | RNA recognition motif (aa654-722) | -18 |
| *Rbl2* | Q64700 | Y632 | RB A (aa414-606) | +26 |
| *Gatad2b* | Q8VHR5 | S123 | p66 CC (158-201) | -35 |
| *Mbd3* | Q9Z2D8 | Y83 | MBDa (79-148) | 0 |
| *Sin3a* | Q60520 | S939 | Sin3a C (887-1187) | 0 |
| *Smarca4* | Q3TKT4 | S1388 | SnAC (1289-1356) | +32 |
| *Med7* | Q9CZB6 | S195 | Med7 (7-164) | +31 |
| *Clock* | O08785 | S91 | HLH (35-85) | +6 |
|  |  | S94 | HLH (35-85) | +9 |
|  |  | S403 | PAS 11 (273-380) | +23 |
|  |  | S406 | PAS 11 (273-380) | +26 |
|  |  | S408 | PAS 11 (273-380) | +28 |
| *Ncl* | P09405 | S605 | RRM1 (574-641) | 0 |
|  |  | S462 | RRM1 (395-460) | +2 |
|  |  | T307 | RRM1 (309-377) | -2 |
|  |  | S563 | RRM1 (489-555) and RRM1 (571-638) | +8 and -8 |
| *Smc1a* | Q9CU62 | S649 | SMC hinge SMC N (512-629) | 0 |

**Supporting information references**
